# Synovial fibroblasts support vascular function in an acute injury-on-a-chip model

**DOI:** 10.1101/2025.07.12.664530

**Authors:** Hannah M. Zlotnick, Declan N. Goddard, Christopher J. Calo, Abhishek P. Dhand, Matthew D. Davidson, Aina Solsona-Pujol, Hannah K. Weppner, Melissa Wong, Carla R. Scanzello, Laurel E. Hind, Jason A. Burdick

## Abstract

Most patients who sustain an acute joint injury develop degenerative joint disease, or osteoarthritis (OA). Animal models have informed the design of OA therapeutics; however, no disease-modifying therapy has successfully translated to human patients. Thus, there is a strong motivation to develop humanized *in vitro* platforms to fill a critical gap in knowledge of disease progression post-injury. Here, we develop an acute injury-on-a-chip model of the synovium, a vascularized, joint-lining tissue that has been implicated in OA progression and as a key driver of joint disease. We apply this chip-based system to investigate crosstalk between endothelial cells, lining an engineered vessel, and synovial fibroblasts, embedded within an extracellular matrix hydrogel. Our data indicate that synovial fibroblasts, rather than initiating disease, attempt to support and maintain vascular function in the presence of acute inflammation (i.e., interleukin-1β). Such knowledge may provide new targets for OA therapeutics, preventing the progression from joint injury to disease in patients.

**Teaser:** In the presence of inflammation, a hallmark of acute injury, synovial fibroblasts work to maintain vascular health.

## Introduction

The incidence of acute joint injuries (e.g., anterior cruciate ligament (ACL) tears) is steadily increasing among young athletes (*1*). This is problematic as the majority (>50%) of patients who sustain a joint injury develop osteoarthritis (OA), a painful, degenerative joint disease, within 20 years (*2*). Currently, the only clinical treatment for end-stage OA is a joint replacement. And, unfortunately, patients under 65 years old who receive a joint replacement have a higher lifetime risk of implant revision (*3*), increasing the opportunities for implant failure and infection. Therefore, there is a strong clinical need to develop new therapies to prevent the progression from joint injury to OA.

There is increased evidence implicating the synovium, a fibrovascular joint-lining tissue, in the progression of joint-wide disease, making tissue-resident synovial cells attractive therapeutic targets to prevent OA (*4*). In fact, synovitis, or synovial inflammation (*5*), is not just an indicator of disease, but may actually cause OA (*6*, *7*). Synovitis is defined by a range of clinical, histological, molecular, and imaging observations (*4*). Notably, the functional capacity of the synovium diminishes with synovitis, whereby lymphatic drainage decreases (*8*, *9*), and immune cells enter the joint via the synovial blood vessels, lined by endothelial cells (*10*). Animal models of acute injury have been instrumental in characterizing synovial pathogenesis (*5*, *11*, *12*); however, within the complex joint environment, it is challenging to examine how specific cell populations and crosstalk between such cells contribute to the initiation and progression of disease.

Tissue-on-a-chip devices and microphysiological systems (MPSs), which often include primary human cells, enable the dissection of cellular interactions due to their modular design. Across the field of musculoskeletal research, MPSs have been developed to model and investigate tendon fibrosis (*13*, *14*), skeletal muscle injury (*15*), articular cartilage disease (*16*), intervertebral disc degeneration (*17*), endochondral ossification (*18*), and synovial disease (*19–21*). For the synovium, current chip-based systems mimic aspects of health and disease, but have primarily focused on therapeutic screening. As a result, these devices maximize throughput, but lack key structural components (e.g., dimensionality) required for mechanistic studies of cellular crosstalk. Namely, existing synovial chips include endothelial cells as either a monolayer, or within a rectangular geometry. While this reduction in tissue structure may be suitable for drug screening, endothelial cell dysfunction has been linked with dimensionality (*22*, *23*). Thus, endothelial cell inclusion within 3D, cylindrical lumens is needed to better understand pathogenesis after acute injury. Recent advances in micromolding and biofabrication have enabled the creation of 3D lumen structures within ECM-based hydrogels (*24–26*). These advances now allow the development of a system of enhanced geometric complexity towards the investigation of the functional response of human synovial fibroblasts and endothelial cells to acute inflammation.

To address these limitations of available platforms of synovial disease in the context of OA, we developed an acute injury-on-a-chip MPS to study synovial fibroblast-endothelial cell interactions and the vascular response of the synovium to acute inflammation *in vitro*. We first motivate the development of this acute injury-on-a-chip MPS through the histological characterization of human synovia from healthy, injured, and diseased human patients. We next utilized traction force microscopy (TFM) to understand how synovial fibroblasts and endothelial cells respond to an inflammatory stimulus, alone and when cultured together on the same substrate. The acute injury-on-a-chip MPS was then developed using a streamlined casting technique with 3D-printed master molds, including endothelialized lumens surrounded by human synovial fibroblasts and acute injury, induced using a pathophysiologic level of the pro-inflammatory cytokine interleukin-1β (IL-1β) (*27*). This model system was then used to measure functional changes in vascular permeability, protein expression, and extracellular matrix (ECM) remodeling over time. By assessing each of these metrics in the presence or absence of synovial fibroblasts, we gain new insights into the distinct role of synovial fibroblasts in maintaining vascular function. Overall, our vascularized acute injury-on-a-chip MPS of the synovial subintima captures key features of this fibrovascular tissue, allowing us to deconstruct complex cell-cell, cell-stimulus, and cell-ECM interactions immediately after a joint injury.

## Results

### Synovial vascularity and cellularity are altered throughout disease progression

Alterations in synovial vascularity and cellularity have been reported throughout the progression from early to end stage OA (*10*). However, it is not fully understood when these changes first occur. To guide the timely delivery of therapeutics aimed at preserving joint function, it is critical to understand this disease progression. Therefore, human synovia were acquired and classified into three groups: healthy (Table S1; organ donor), injured (Table S2; meniscus arthroscopy), and diseased (Table S3; total knee arthroplasty). All synovia were histologically sectioned and stained with hematoxylin and eosin to assess both vascularity and cellularity (Fig. 1A, B). The healthy samples showed a defined intima, around one cell layer thick, and sparsely vascularized subintima, whereas the injured and diseased tissues appeared markedly different than the healthy tissue (Fig. 1C). To better understand these differences, four blinded graders scored the sections across six grading criteria: vascularity, subsynovial infiltrates, lining hyperplasia, fibrosis, vasculopathy, and perivascular edema. Both injured and diseased synovial tissues showed a significant increase in vascularity (Fig. 1D), defined by a greater number of vessels in the field of view. Accompanying this increase in vasculature, there were significantly more subsynovial infiltrates, denoting immune cell populations, in both the injured and diseased synovial tissues (Fig. 1E). There was no significant consensus on differences in the lining hyperplasia (Fig. 1F), likely due to the presence of lining erosion in some of the tissues, a sign of synovial tissue damage that presents as a decrease in cellularity at the intimal boundary that is opposite that of lining hyperplasia (*10*). Additional scoring criteria, including fibrosis, vasculopathy, and perivascular edema, did not show differences between groups (Fig. S1). For all scoring criteria, the intraclass correlation coefficient (ICC) values were greater than 0.6, excluding vasculopathy, indicating sufficient grader-to-grader agreement.

**Figure 1.**
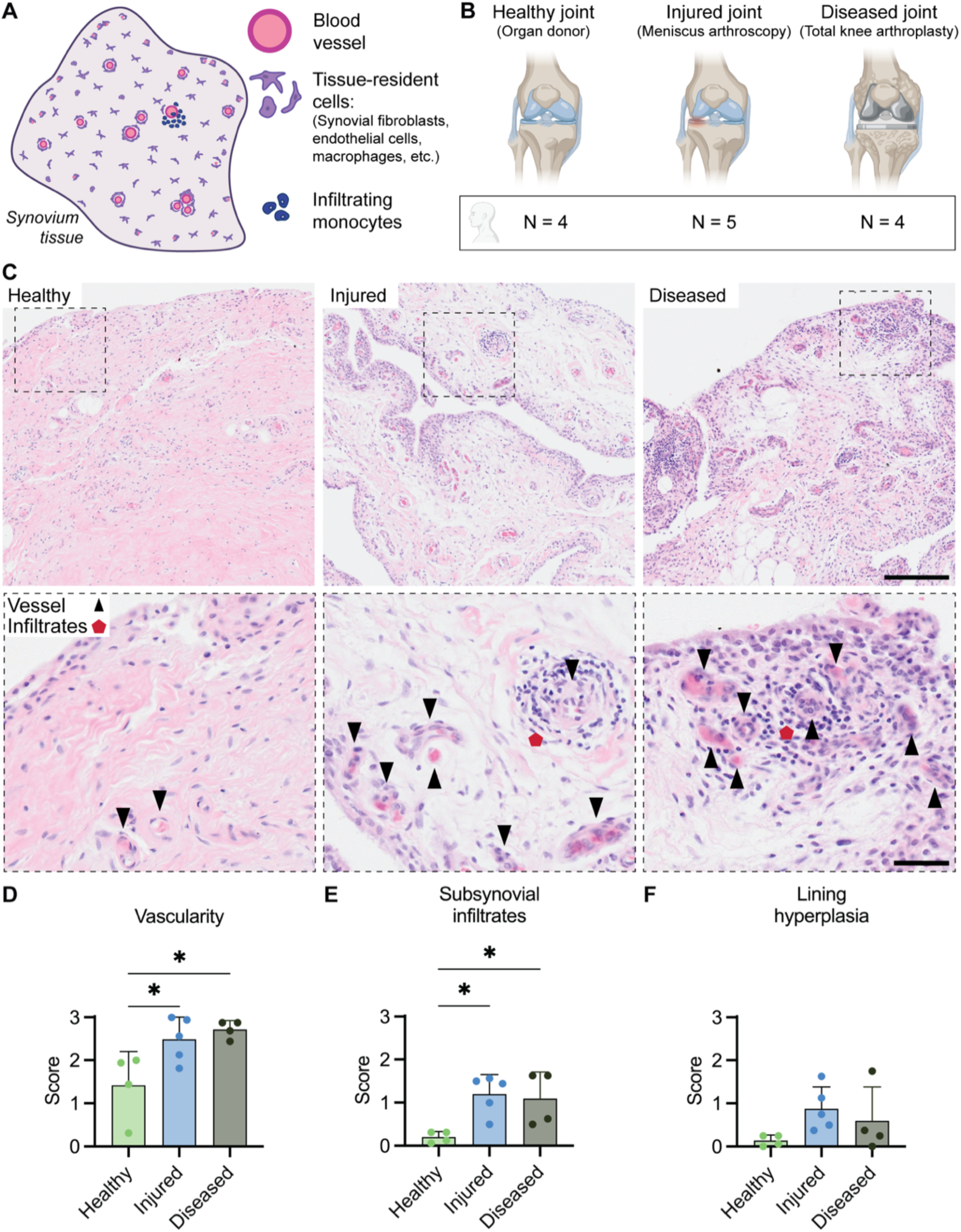
Knee injury increases vascularity and subsynovial infiltrates in the human synovium. (A) Schematic of synovium stained with hematoxylin and eosin, denoting blood vessels (pink, surrounded by purple nuclei) and infiltrating immune cell populations (deep purple, rounded) amidst tissue-resident cells. (B) Overview of human donor synovia acquired from healthy (n = 4), injured (knee meniscus injury; n = 5), and diseased (total knee replacement; n = 4) knees. (C) Hematoxylin and eosin staining of donor tissues. Scale bars = 200 µm (top), 50 µm (bottom). (D-F) Histological scoring for vascularity, subsynovial infiltrates, and lining hyperplasia. For each scoring criterion, scores ranged from 0 (physiologic) to 3 (pathophysiologic). n = 4 trained, blinded scorers. Data plotted as the average score for each tissue section by 4 scorers and bars represent mean + std dev. ANOVA with Tukey post-hoc comparisons. \**P* < 0.05.

Taken together, these data highlight how joint injuries, such as meniscal tears, disrupt the synovial environment, leading to vascular changes that can be observed at the time of surgical arthroscopy for meniscal repair in human patients. Surgical arthroscopy may occur anywhere from 1-week to several years after an initial injury. This is often the earliest timeframe where synovia explants can be acquired. Therefore, it is unknown how the endothelial cells and the surrounding stromal populations immediately respond to acute injury. Human cell-based models offer an alternative for investigation into this early timeframe post-injury.

### Synovial fibroblasts play a supportive role in maintaining endothelial cell traction

After an injury, changes in the synovial environment eventually result in an increase in vasculature and immune cell infiltrates. It is unknown what synovial cell populations or molecular cues drive these changes. To investigate how local cell populations within synovial tissue respond to acute injury, we utilized traction force microscopy (TFM). TFM is an established tool in the field of mechanobiology that enables the quantification of cellular traction forces in a controlled system that relies on single cells interacting with their substrate biophysically and in the presence of biochemical factors within the culture media. To assess cell-generated traction forces, both endothelial cells and synovial fibroblasts were individually seeded on polyacrylamide (PA) gels and then in co-culture for TFM analysis.

To test the impact of acute injury on endothelial cell traction, human umbilical vein endothelial cells (HUVECs) were seeded on PA gels in the presence or absence of IL-1β, mimicking acute injury (Fig. 2A). Confocal images were acquired before (Fig. 2B) and after cell lysis, to track the displacement of the fiducial markers. The greatest bead displacements and traction stresses were observed at the cell periphery (Fig. 2C, D). A similar protocol was carried out for the synovial fibroblasts to determine the influence of IL-1β on synovial fibroblast traction stress (Fig. 2E-H). While TFM is traditionally performed by seeding a single cell type on a 2D gel, we also utilized this assay to determine the effect of one cell population on another by seeding both HUVECs and synovial fibroblasts on the same gel, but not in physical contact with each other, and separately analyzing each population (Fig. 2I-L).

**Figure 2.**
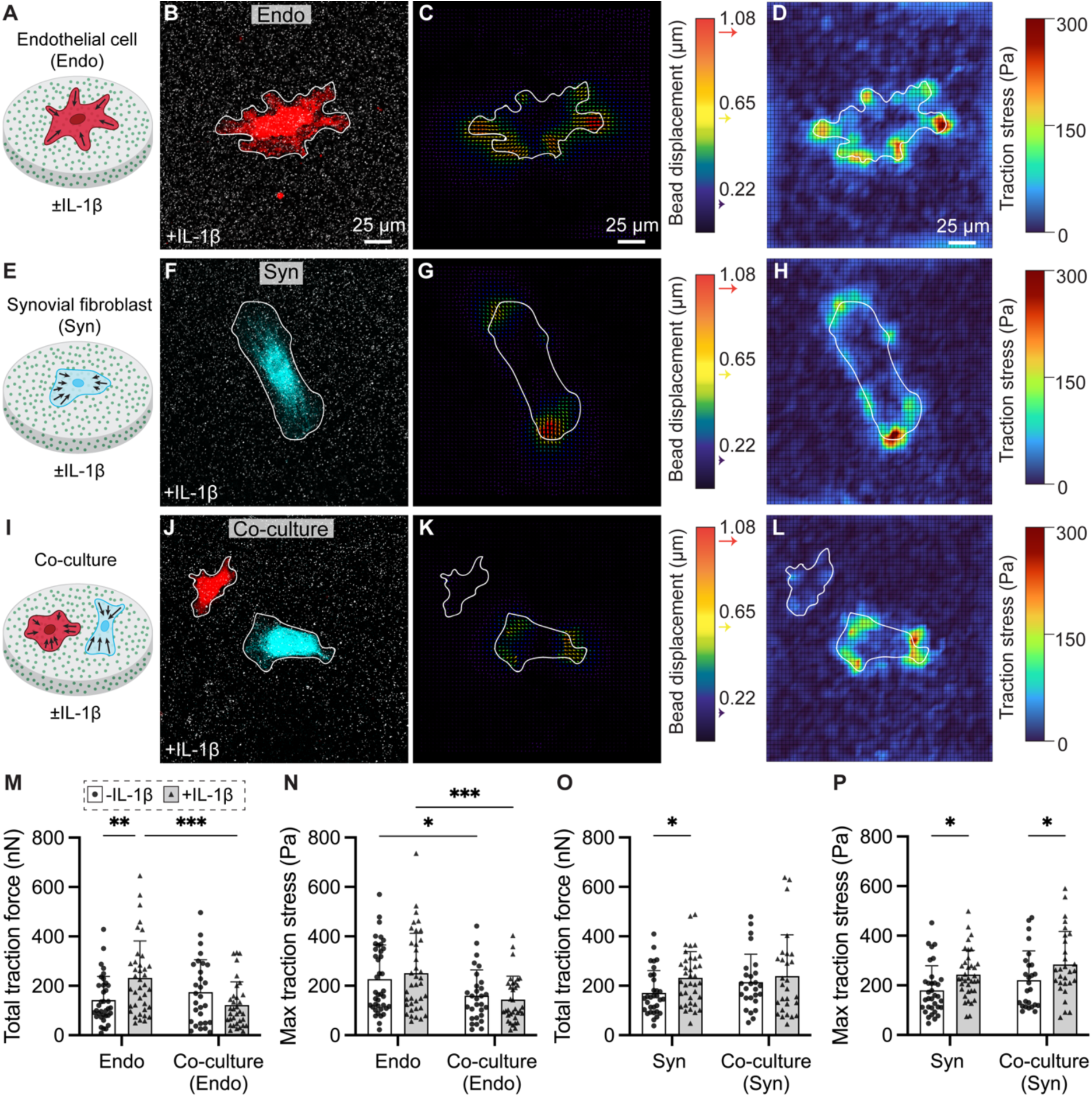
Human synovial fibroblasts reduce endothelial cell traction in the presence of an inflammatory factor. Traction force microscopy (TFM) was conducted for cells (Red fluorescent protein (RFP)-human umbilical vein endothelial cells (HUVECs) alone, human synovial fibroblasts alone, co-cultures of both HUVECs and synovial fibroblasts) seeded onto polyacrylamide (PA) gels containing embedded fluorescent beads either without (-) or with (+) IL-1β for 24 h. (A) Schematic of TFM for RFP-HUVECs alone, as well as representative (B) fluorescent image of RFP-HUVEC before cell lysis, (C) vector field (bead displacement) from a single endothelial cell, post-lysis calculated via particle image velocimetry (PIV) analysis, and (D) corresponding heat map of the traction stress for an isolated endothelial cell, calculated via Fourier transform traction cytometry (FTTC) analysis. (E) Schematic of TFM for human synovial fibroblasts alone, as well as representative (F) fluorescent image of human synovial fibroblast (cell tracker, cyan) before cell lysis, (G) bead displacement vector field after cell lysis, and (H) traction stress heat map. (I) Schematic of TFM for co-culture of RFP-HUVECs (red) and human synovial fibroblasts (cyan), as well as (J) fluorescent images of RFP-HUVEC and synovial fibroblast on the same gel, in proximity to each other, but not touching, (K) bead displacement vector field after cell lysis, and (L) traction stress heat map for both cells. (M, N) Total traction force and max traction stress for the endothelial cells, in isolation and co-culture (with synovial fibroblasts). White bars = cultured in EGM-2, grey bars = cultured in EGM-2 + 100 pg mL^-1^ IL-1β. n = 29-40 cells/group. 2-way ANOVA with Fisher’s LSD test. Bars represent mean + std dev. *P*\* < 0.05, *P*\*\* < 0.01, *P*\*\*\* < 0.001. (O, P) Total traction force and max traction stress for the synovial fibroblasts, in isolation and co-culture (with endothelial cells). White bars = cultured in EGM-2, grey bars = cultured in EGM-2 + 100 pg mL^-1^ IL-1β. n = 26-37 cells/group. Bars represent mean + std dev. 2-way ANOVA with Fisher’s LSD test. \**P*< 0.05.

The low dose of 100 pg mL^-1^ IL-1β, applied for only 24 hours, induced a disease-like state in the endothelial cells, increasing total traction force (Fig. 2M), while maintaining cell area (Fig. S2). However, when the endothelial cells were cultured with synovial fibroblasts, creating a more physiologic environment, their total traction force significantly decreased in the presence of IL-1β. This suggests that synovial fibroblasts have the ability to regulate endothelial cell traction. In alignment with the total traction force data for the endothelial cells, we also observed that the maximum traction stress of the endothelial cells decreased with the addition of synovial fibroblasts (± IL-1β) (Fig. 2N), again suggesting a supportive role for the synovial fibroblasts. The endothelial cells did not appear to influence the synovial fibroblasts in a reciprocal fashion (Fig. 2O, P), as synovial fibroblast contractility was largely unaffected by the presence of the endothelial cells. However, similar to the endothelial cells, the synovial fibroblasts displayed enhanced contractility with the addition of IL-1β.

Overall, IL-1β increased cellular traction for both the endothelial cells and synovial fibroblasts. When these cells were cultured together to mimic the local cellular environment in the synovial subintima, we discovered that the synovial fibroblasts unexpectedly decreased endothelial cell traction. To date, most studies have focused on the pro-inflammatory, or pro-fibrotic role of synovial fibroblasts (*28*). Our findings support the further investigation of cellular crosstalk in the synovium, particularly through the use of multi-cellular models.

### Development of a vascularized acute injury-on-a-chip model

To further investigate endothelial cell-synovial fibroblast crosstalk after an acute injury, we developed a vascularized acute injury-on-a-chip model of the synovial subintima. These chip systems were fabricated, similar to previous studies (*10*, *11*), by inserting a polydimethylsiloxane (PDMS) rod between two PDMS layers (Fig. 3A). However, instead of using photolithography-based master molds to generate the PDMS layers, we designed durable, heat-resistant 3D-printed master molds (Fig. S3).

**Figure 3.**
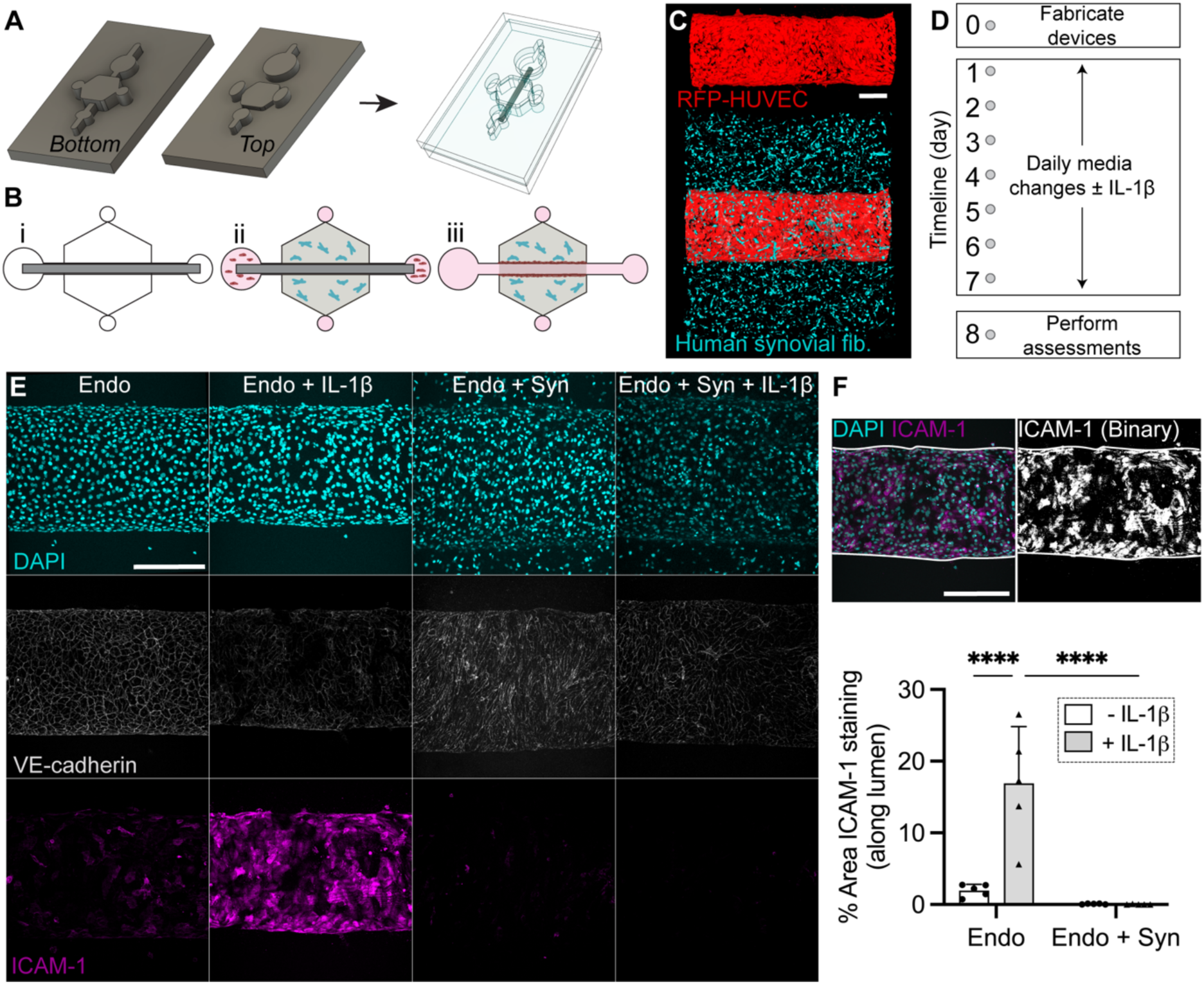
Synovial fibroblasts reduce ICAM-1 expression in an acute injury-on-a-chip model system. (A) Computer aided design (CAD) rendering and bi-layered polydimethylsiloxane (PDMS) microfluidic devices. (B) Schematic illustrating the stepwise assembly of the microfluidic acute injury-on-a-chip system, including PDMS casting from the master mold (i), collagen polymerization (± human synovial fibroblasts) (ii), and endothelial cell seeding after removal of the inner rod (shown in dark grey) (iii). (C) 3D reconstruction of acute injury-on-a-chip system containing endothelial cells (red; RFP-HUVECs) within the lumen (top) and both synovial fibroblasts (cyan; cell tracker labeled) within the collagen gel and endothelial cells within the lumen (bottom). Scale bar = 200 µm. (D) Timeline for the induction of acute injury via the administration of daily IL-1β (100 pg mL^-1^). (E) Immunofluorescence staining (DAPI—cyan (top), VE-cadherin—white (middle), ICAM—magenta (bottom)), of devices at day 8. Endo: endothelialized lumen cultured in EGM-2. Endo + IL-1β: endothelialized lumen cultured in EGM-2 + IL-1β. Endo + Syn: endothelialized lumen surrounded by human synovial fibroblasts (in the collagen gel) cultured in EGM-2. Endo + Syn + IL-1β: endothelialized lumen surrounded by human synovial fibroblasts (in the collagen gel) cultured in EGM-2 + IL-1β. n = 5 lumens/group. Scale bar = 200 µm. (F) Quantification of ICAM-1 staining from binarized max-projection images (n = 5 lumens/group). Scale bar = 200 µm. White bars = cultured in EGM-2, grey bars = cultured in EGM-2 + 100 pg mL^-1^ IL-1β. Bars represent mean + std dev. 2-way ANOVA with Fisher’s LSD test. \*\*\*\**P*< 0.0001.

Using the assembled PDMS devices, we formed the cell-based lumens by first injecting a collagen solution (either without or with synovial fibroblasts) into the top port of the devices and allowing the collagen to polymerize. Next, we injected endothelial cells into the side ports of the devices (Fig. 3B) and removed the inner PDMS rod. Upon rotation of the devices after seeding, endothelial cells coated the circumference of the lumens. Additionally, due to the bi-layered nature of the devices, synovial fibroblasts fully surrounded the lumens in the outer gel (Fig. 3C). To repeatably induce acute injury, we dosed the lumens with 100 pg mL^-1^ of IL-1β, a pro-inflammatory cytokine, daily through day 8 of culture (Fig. 3D). We intentionally selected this dosing scheme to mimic the pathologic environment of the human knee joint after an acute injury. The detected range of IL-1β in the human knee post-injury is 0.3 to 135.2 pg mL^-1^, as quantified from synovial fluid aspiration (*27*).

After 8 days of culture, we stained the lumens to assess cellularity (DAPI), endothelial cell contacts (VE-cadherin), and vascular inflammation (ICAM-1) in the presence and absence of both synovial fibroblasts and IL-1β (Fig. 3E). From the DAPI images, we clearly observed cells in the extraluminal space of the synovial fibroblast-seeded groups, but not the endothelial cell-alone groups (Endo, Endo + IL-1β), surrounded by acellular collagen. VE-cadherin staining, denoting cell-cell junctions between the endothelial cells, indicated complete vascular formation across all groups. Most interestingly, we noted elevated ICAM-1 staining in the Endo + IL-1β devices, compared to all other groups. Upon quantification of ICAM-1 staining, IL-1β significantly increased endothelial inflammation, and the addition of synovial fibroblasts significantly lowered the expression of this marker even in the presence of IL-1β (Fig. 3F). This again suggests that synovial fibroblasts play a supportive role in maintaining vascular function, particularly in the presence of inflammation.

### Synovial fibroblasts enhance vascular barrier function

Having successfully recreated tubular, endothelialized vessels, we applied this model system to investigate the influence of synovial fibroblasts (± IL-1β) on vascular barrier function. While vascular barrier function can be quantified within the knee joint of animal species (*29*), it is challenging, if at all possible, to pinpoint how specific cell populations influence vascular permeability *in vivo*. MPSs, made from optically transparent elastomers, enable live imaging of fluorescent dye diffusion (Fig. 4A), and quantification of diffusive permeability (*P*_d_) (*24*).

**Figure 4.**
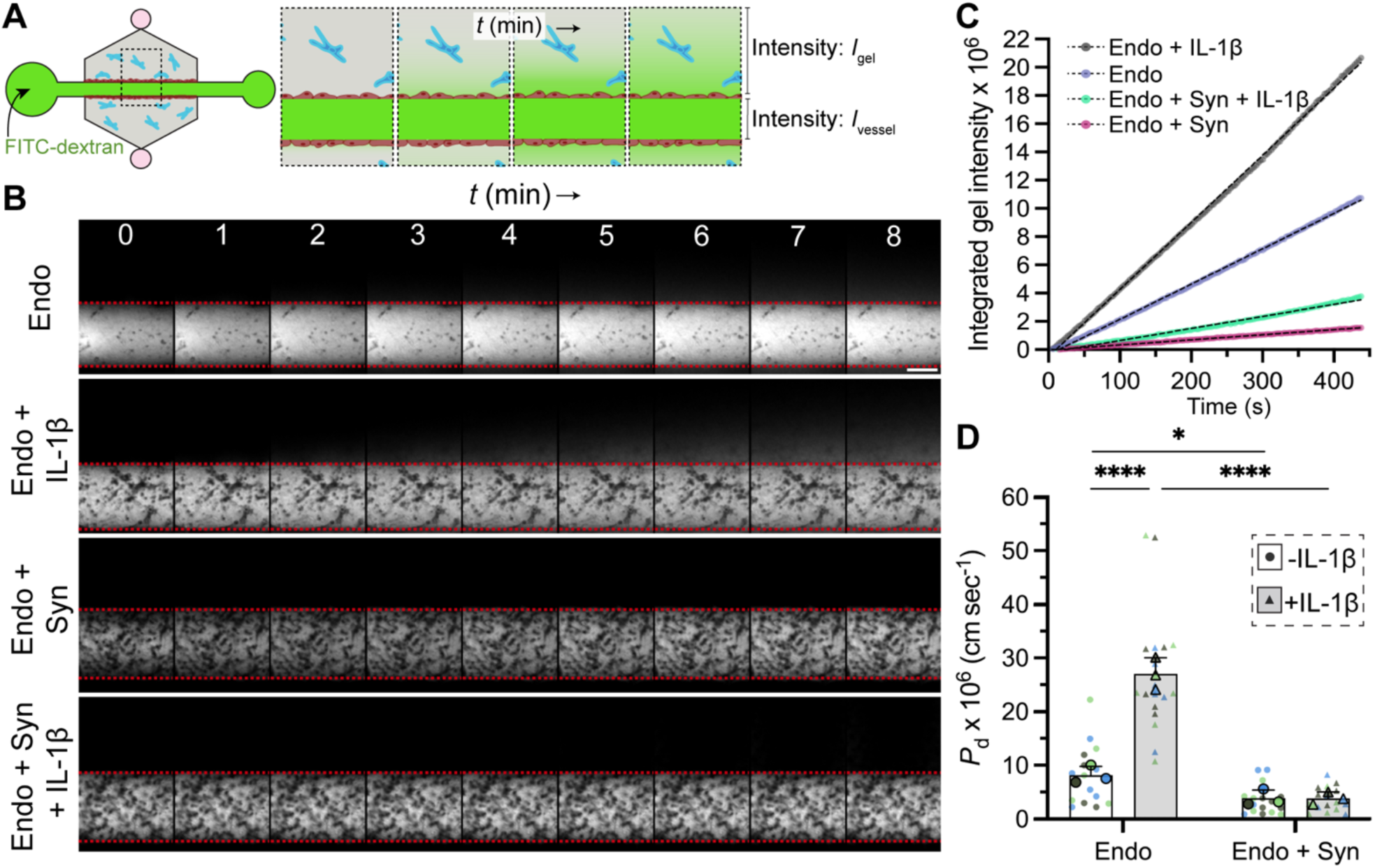
Synovial fibroblasts enhance vascular barrier function in an acute injury-on-a-chip model system. (A) Schematic of vascular permeability assay. FITC-Dextran (greyscale) diffusion out of the vessel was imaged over time. The *I*_gel_ and *I*_vessel_ were calculated in MATLAB, enabling quantification of diffusive permeability (*P*_d_). (B) Representative time-lapse microscopy images over 8 min. Endo: endothelialized lumen cultured in EGM-2. Endo + IL-1β: endothelialized lumen cultured in EGM-2 + IL-1β. Endo + Syn: endothelialized lumen surrounded by human synovial fibroblasts (in the collagen gel) cultured in EGM-2. Endo + Syn + IL-1β: endothelialized lumen surrounded by human synovial fibroblasts (in the collagen gel) cultured in EGM-2 + IL-1β. The dashed red line denotes the edge of the vessel. Scale bar = 200 µm. (C) Representative integrated gel intensity plot over time (colored data) and simple linear regression for each group (dashed lines). (D) *P*_d_ of each group. Colored replicates (n = 4-6 lumens/group/donor). The mean for each biological replicate is marked by the larger data point. Bars represent overall mean + std dev. 2-way ANOVA with Fisher’s LSD test. \**P* < 0.05. \*\*\*\**P* < 0.0001.

To measure vascular barrier function within our acute injury-on-a-chip, we injected fluorescein isothiocyanate (FITC)-dextran (10 kDa) into each lumen, and acquired images every 3 s for 8 min (Fig. 4B). The fluorescent intensity of the gel was plotted over time to visualize FITC-dextran accumulation outside of the vessel (Fig. 4C), and diffusive permeability was calculated as previously reported (*24*). When first comparing the Endo and Endo + IL-1β groups, the addition of IL-1β significantly increased vascular leakiness (Fig. 4D). For reference, the average diffusive permeability for the Endo + IL-1β group overlapped with the diffusive permeability of the acellular collagen gel at the same collagen concentration (Fig. S4). Therefore, even the low dosage of IL-1β used in this study elicited a drastic response in the endothelial cell-alone group. Moving forward, we added synovial fibroblasts into the chip to first assess how they contribute to maintaining vascular homeostasis within the synovium, by comparing the Endo and Endo + Syn groups. Here, we observed improved barrier function with the addition of synovial fibroblasts to the collagen gel. The addition of synovial fibroblasts had an even more pronounced impact on the IL-1β groups, where the addition of synovial fibroblasts resulted in maintenance of vascular barrier function in the presence of IL-1β. It is important to note that the addition of synovial fibroblasts to the collagen gel, in the absence of endothelial cells, did not influence barrier function, in comparison to acellular collagen (Fig. S4). Therefore, when comparing the Endo + IL-1β and Endo + Syn + IL-1β groups, the enhanced barrier function in the Endo + Syn + IL-1β group is likely due to cell-cell crosstalk between the endothelial cells and synovial fibroblasts.

### Synovial fibroblasts downregulate disease signals

In the presence of acute inflammation (i.e., IL-1β treatment), the cells within the synovium differentially respond to either subdue or amplify disease. To deconstruct the influence of both endothelial cells and synovial fibroblasts, we collected the culture media from the devices on day 8 for analyte analysis using a Luminex multiplex bead-based enzyme-linked immunosorbent assay (Fig. 5A). The profiled analytes included both pro- and anti-inflammatory mediators. Our results revealed two primary clusters of protein expression, one dominated by the presence of synovial fibroblasts, and the other governed by the addition of IL-1β. Interleukin-8 (IL-8), monocyte chemoattractant protein-1 (MCP-1), and interleukin-1α (IL-1α) appeared to be downregulated by the addition of synovial fibroblasts, in comparison to the Endo control (Fig. 5B). Indeed, both MCP-1, a chemokine, and IL-1α, a pro-inflammatory cytokine, significantly decreased with the addition of synovial fibroblasts, in the presence of IL-1β (Fig. 5C, D). This suggests that the synovial fibroblasts again reduced disease in the acute injury-on-a-chip MPS.

**Figure 5.**
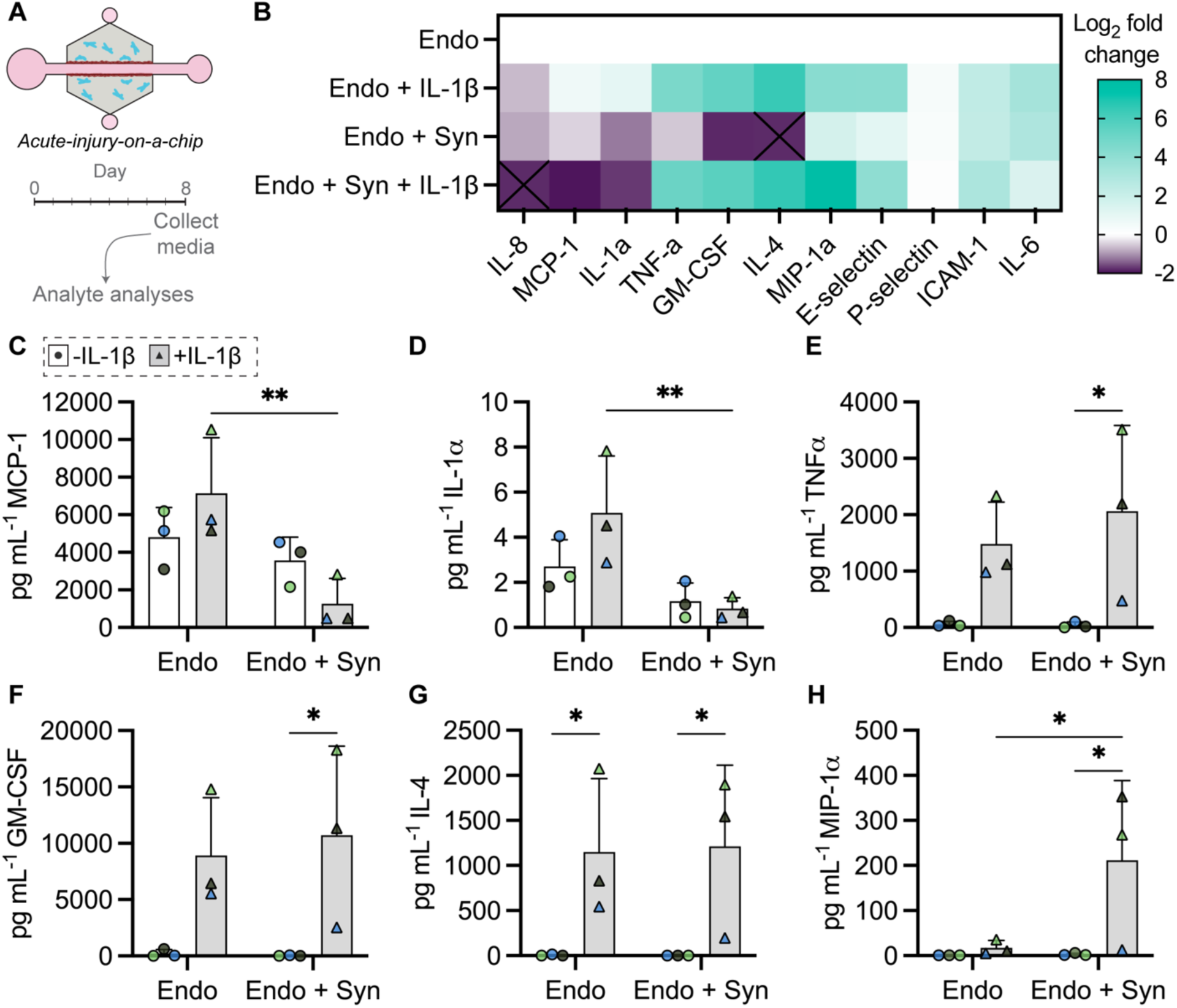
Synovial fibroblasts downregulate MCP-1 and IL-1α in the presence of acute inflammation. (A) Schematic of media collection from acute injury-on-a-chip devices for analyte analyses via a multiplex human inflammation panel. (B) Secreted protein expression, normalized to the Endo group (Log_2_ fold change). Endo: endothelialized lumen cultured in EGM-2. Endo + IL-1β: endothelialized lumen cultured in EGM-2 + IL-1β. Endo + Syn: endothelialized lumen surrounded by human synovial fibroblasts (in the collagen gel) cultured in EGM-2. Endo + Syn + IL-1β: endothelialized lumen surrounded by human synovial fibroblasts (in the collagen gel) cultured in EGM-2 + IL-1β. Upregulated: teal, downregulated: purple, in respect to the Endo group. Black ‘X’ denotes no detectable signal. No signal was detected for IFN-α, IFN-γ, IL-1β, MIP-1β, IL-10, IL-13, IL-12p70, IL-17A, IP-10 across all groups. (C-H) Concentrations (pg mL^-1^) of secreted proteins that showed statistical significance. Colored replicates (n = 6 lumens pooled/group/donor). Bars represent overall mean + std dev. 2-way ANOVA with Fishers LSD test. \**P* < 0.05.

For the following soluble proteins: tumor necrosis factor-α (TNF-α), granulocyte-macrophage colony-stimulating factor (GM-CSF), interleukin-4 (IL-4), macrophage inflammatory protein-1α (MIP-1α), and E-selectin, IL-1β treatment predictably increased their expression. Notably, TNF-α, GM-CSF, IL-4, and MIP-1α were significantly elevated in the Endo + Syn + IL-1β group, in comparison to the Endo + Syn group (Fig. 5E-H). Hence, IL-1β treatment stimulated the production of the pro-inflammatory cytokines, TNF-α and GM-CSF, as well as the anti-inflammatory cytokine, IL-4, and inflammatory chemokine, MIP-1α. Because these factors appear elevated in the Endo + IL-1β group as well as the Endo + Syn + IL-1β group, their expression is likely from the endothelial cells. Other analytes including P-selectin, ICAM-1, and interleukin-6 (IL-6) showed no differences between groups (Fig. S5). Overall, this analysis suggests that, within our model system, the synovial fibroblasts work to maintain homeostasis while the endothelial cells, in the presence of IL-1β, upregulate inflammation.

### Acute inflammation disrupts cell-ECM interactions

Within the native synovium, there is an increase in blood vasculature and immune cell infiltrates with OA disease progression (*10*). For new blood vessels to form, the existing endothelial cells must receive angiogenic cues. The endothelial cells must also secrete enzymes (i.e., matrix metalloproteinases—MMPs) to breakdown the vascular basement membrane, remodel the perivascular ECM, and migrate to form new vasculature (*30*). Within our acute injury-on-a-chip, we can track perivascular ECM remodeling via the inclusion of fluorescently labeled collagen in the gel surrounding the lumen. And so, we fabricated lumens surrounded by fluorescently labeled collagen, added at the time of cell seeding, and assessed collagen remodeling at the day 8 timepoint.

The 3D images of fluorescent collagen organization revealed a striking layer of dense collagen that developed directly around the lumens (Fig. 6A, B), highlighting the importance of using biomaterials that enable cells to self-assemble 3D structures. The endothelial cells appeared to drive this collagen rearrangement, as the thin barrier was present in the Endo group, but not the acellular or synovial fibroblast alone conditions (Fig. S6). The addition of IL-1β significantly disrupted the collagen layer surrounding the endothelialized lumens (Fig. 6C). In fact, the Endo + IL-1β group appeared very similar to the acellular collagen control without any endothelial cells. When synovial fibroblasts were added to the outer gel, the collagen layer around the lumen dramatically increased. Because endothelial cells, and not synovial fibroblasts, appear to drive this remodeling, we believe that cell-cell crosstalk enhances endothelial cell-ECM remodeling in the presence of synovial fibroblasts. Lastly, the addition of IL-1β to the Syn + Endo group reduced this collagen remodeling, creating a more patchy-like structure, although more visible than the Endo + IL-1β group. Overall, these results highlight one of the many strengths of this 3D model system. It would be challenging, if at all possible, to assess physiologic ECM remodeling in an endothelial monolayer or within tissue sections.

**Figure 6.**
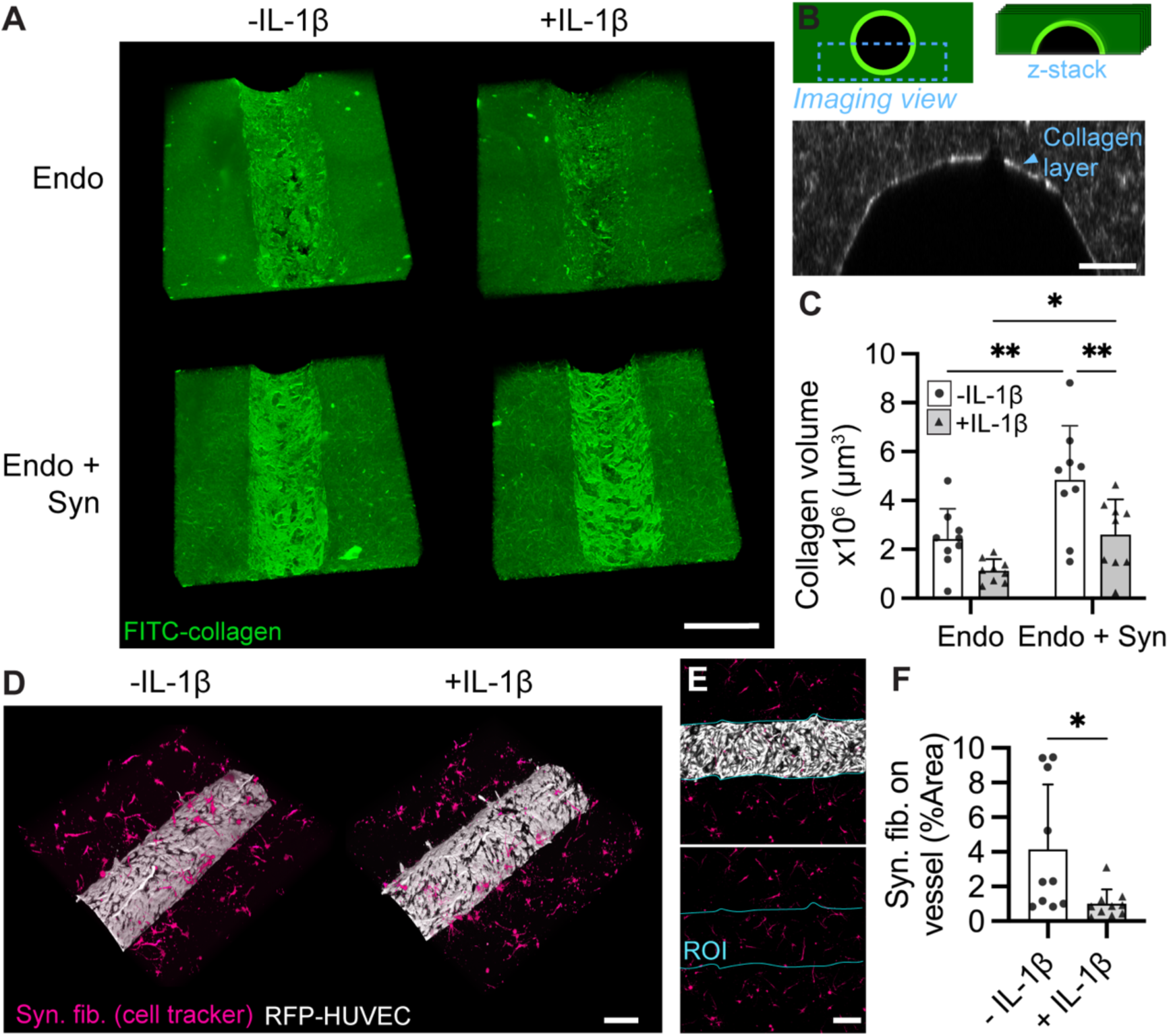
Acute inflammation alters extracellular matrix structure and fibroblast localization near vasculature. (A) Representative 3D reconstructions from each group, highlighting the denser collagen layer (or lack thereof) along the lumens that developed over time. Images were taken at day 8, and Fluorescein isothiocyanate (FITC)-collagen was added at the time of cell seeding. Endo: endothelialized lumen cultured in EGM-2. Endo + IL-1β: endothelialized lumen cultured in EGM-2 + IL-1β. Endo + Syn: endothelialized lumen surrounded by human synovial fibroblasts (in the collagen gel) cultured in EGM-2. Endo + Syn + IL-1β: endothelialized lumen surrounded by human synovial fibroblasts (in the collagen gel) cultured in EGM-2 + IL-1β. Scale bar = 400 µm. (B) Schematic of imaging pipeline and collagen layer identification. Scale bar = 100 µm. (C) Volumetric quantification of collagen volume along the vessel. n = 9 lumens/group. Bars represent mean + std dev. 2-way ANOVA with Fisher’s LSD test. \**P* < 0.05. \*\**P* < 0.01. (D) Representative 3D reconstructions of the coculture groups (Endo + Syn) ± IL-1β. Synovial fibroblasts (pink), and Red fluorescent protein (RFP)-human umbilical vein endothelial cells (HUVECs) (white). Scale bar = 100 µm. (E) Max projection image showing region of interest (ROI) for quantitative analysis. The signal from the synovial fibroblasts was quantified within the vessel region. Scale bar = 200 µm. (F) Synovial fibroblast localization on the vessel, reported as %Area. n = 10 lumens/group. Bars represent mean + std dev. Unpaired t-test. *P < 0.05.

In conjunction with our analysis of cell-ECM remodeling, we assessed the localization of the synovial fibroblasts within the acute injury-on-a-chip model. Here, we imaged both the synovial fibroblasts and the endothelial cells to determine how IL-1β influenced synovial fibroblast localization along the lumen. Representative 3D images highlight an increase in synovial fibroblasts conforming to the curved vessel geometry in the absence of IL-1β (Fig. 6D). To quantify this finding, a region of interest was established including the complete vessel area (Fig. 6E). Significantly more synovial fibroblasts were localized along the vessel in the absence of IL-1β. The low dose of daily IL-1β treatment impacted the localization of the synovial fibroblasts, driving them further from the vessels. Therefore, across all other outcome measures (i.e., traction force microscopy, ICAM-1 staining, vascular permeability, soluble analyte analysis, collagen remodeling), the observed positive impact of the synovial fibroblasts may not be explained by physical proximity, or attachment, to the endothelial cells. This suggests that cell-cell crosstalk, through the release of soluble signals from the synovial fibroblasts, may modulate vascular homeostasis.

## Discussion

The development of effective therapies for both synovitis and OA requires an increased understanding of the complex joint environment, both in homeostasis and throughout disease progression. Characterization of the human synovia throughout health, injury (i.e., meniscus tear), and disease (i.e., OA) using patient explants reveals significant changes in the synovial subintima after injury, where there is an increase in blood vessels and immune cell infiltrates. Limited by the lack of specific time points prior to injury provided by these synovia explants, we sought to develop human cell-based models to investigate early cell-cell crosstalk between endothelial cells and synovial fibroblasts in the synovium after an acute injury. This study both develops and utilizes an acute injury-on-a-chip model of the synovial subintima to dissect these cellular behaviors and provides a never-before-seen lens into the response of both endothelial cells and synovial fibroblasts to acute inflammation.

Increased traction stresses are observed for endothelial cells and synovial fibroblasts in response to IL-1β stimulation, a pro-inflammatory stimulus. When both cell populations are seeded on the same substrates, a surprising influence of synovial fibroblasts on endothelial cell traction in the presence of IL-1β is observed. In fact, synovial fibroblasts return endothelial cell traction back to baseline. This result motivates the development of an acute injury-on-a-chip MPS to further investigate endothelial cell-synovial fibroblast crosstalk. 3D-printed master molds enable the production of a bi-layered device with an inner hollow lumen that could be seeded with endothelial cells, surrounded by an extracellular matrix containing synovial fibroblasts. Across multiple outcome measures (i.e., ICAM-1 staining, vascular permeability, protein expression, ECM remodeling), the presence of synovial fibroblasts reduces a disease-like vascular phenotype within the model.

Our data suggests that synovial fibroblasts, or a sub-population thereof, have the functional capacity to support vascular function. While numerous studies have focused on the deleterious role of synovial fibroblasts in OA pathogenesis (*28*, *31*), this work offers a novel role for these cells. Interestingly, when co-culturing synovial fibroblasts with endothelial cells, they can assume mural cell and perivascular phenotypes, as identified with single-cell RNA-sequencing (*32*). However, when cultured alone, synovial fibroblasts do not assume these identities. This suggests that the majority of *in vitro* studies investigating synovial fibroblasts in the absence of endothelial cells may miss the functional capacity of these cells to stabilize and maintain vascular homeostasis. Given our results, an effective therapeutic for synovitis may target synovial fibroblasts. While it is unknown how the vessels within the synovium become disrupted over the course of time, we know from histology that vascular changes occur. Potentially, the synovial fibroblasts, mural cell populations, or a subset thereof, lose their ability to stabilize the vasculature, and this could be targeted via therapeutic delivery. Targeting may occur through antibody-drug conjugates. Single-cell RNA-sequencing of synovial fibroblasts has uncovered multiple distinct cell markers (*11*, *33*). Using these markers, antibodies may be designed to specifically target synovial subpopulations.

Every MPS has limitations in that it can never fully recapitulate all aspects of the *in vivo* environment. Adding too much complexity to a MPS reduces the throughput and usability. And so, care must be taken to intentionally build complexity into a MPS to answer a specific research question, or effectively screen therapeutics (*34*). Therefore, within our acute injury-on-a-chip model system we incorporated the key aspects of synovial physiology that were critical to understand how endothelial cells and synovial fibroblasts contribute to human synovial homeostasis and disease progression. To induce “injury,” we utilize a pathophysiologic dose of IL-1β observed in human knees after acute injury (*27*). In other synovium-on-a-chip studies, researchers have employed much higher dosages of pro-inflammatory cytokines that are 10-1000x outside this range. We also create a 3D chip design, allowing synovial fibroblasts to surround a tubular vessel, similar to the structure of the synovial subintima. This dimensionality is unique, and differs from the majority of MPS studies utilizing thin, relatively 2D channels.

There are further complexities that could be added to our system. Namely, tissue-resident synovial macrophages, both lining and interstitial, are important cell populations within the synovium that constantly survey their local tissue environment and can promote the resolution or activation of inflammation (*35–37*). While these cells are not a focus of our study, tissue-resident synovial macrophages could be integrated into this chip design to further investigate how a tertiary cell population influences endothelial cells and synovial fibroblasts. Another area for further study would be the integration of active fluid flow via a peristaltic pump into the chip design. To circumvent the need for active fluid flow, we changed the media on these devices daily. Also, due to the size of the media ports, there was a pressure differential in the devices leading to gravity-driven flow upon injection of media. This led to excellent vascular barrier function within devices, on par with that reported for devices with active fluid flow (*24*). Lastly, after an injury, the synovial joint environment is complex, involving the upregulation of multiple soluble factors. While we reduce this multifaceted environment to a single cytokine, IL-1β, the inclusion of any number of cytokines to mimic features of the joint environment or the use of patient synovial fluid with acute injury could improve model relevance. Overall, the modularity of MPSs is really an advantage, not a limitation. Different cell populations, external cues, or soluble factors can be added or subtracted from the system in a methodical manner, leading to new discoveries that would be impossible in any *in vivo* system.

The primary objective of this report is to elucidate the early response of endothelial cells and synovial fibroblasts to acute injury, which is extremely challenging, if possible, to do *in vivo*. By developing novel human cell-based culture systems, we uncover a new role of synovial fibroblasts in supporting vascular function. While established to study acute injury, the developed MPS could be adapted to investigate late-stage OA, as well as other synovial diseases, such as rheumatoid arthritis (RA) (*38*). Because vascular transport is critical to joint homeostasis, there are numerous applications of this 3D, vascularized synovium model. Most excitingly, our data suggest that synovial fibroblasts may be powerful targets for therapeutic delivery post-injury to preserve joint function and prevent synovitis.

## Materials and Methods

### Histological analyses and scoring

Deidentified human synovia from organ donors (Table S1), patients undergoing meniscal arthroscopy (Table S2), or total knee arthroplasty (Table S3) were previously banked at Rush University Medical Center and obtained from IRB-approved repositories (ORA 08082803 and ORA 00011021). Samples were taken from the suprapatellar region of the knee, formalin-fixed, dehydrated, and paraffin-embedded for histological sectioning (7 µm). Tissue sections were stained with hematoxylin (26030-20, Electron Microscopy Sciences) and eosin (HT110116, Millipore Sigma), and imaged using a slide scanner (Evident Slideview VS200).

Four blinded, trained reviewers scored the hematoxylin and eosin synovia sections for signs of inflammation (sublining infiltrate, vascularity, lining hyperplasia), and damage (fibrosis, vasculopathy, perivascular edema) using an integer scale (0–3), with 0: physiologic and 3: severely pathologic, respective to each category (*10*, *39*). Average scores were plotted in Figure 1 ± standard deviation. The intraclass correlation (ICC) (Two-way, mixed-effects model for consistency, single measurement) for each metric was calculated using a custom MATLAB script. ICC values were > 0.60, apart from vasculopathy (ICC: 0.58), indicating good inter-rater reliability (Table S4).

### Traction force microscopy (TFM)

Polyacrylamide (PA) hydrogels (5 kPa) were polymerized in methacrylated glass well-bottom plates. Briefly, an Acrylamide: Bisacrylamide (5089990118, Fisher Scientific) solution was diluted (1:5) in distilled water (DI) with ammonium persulfate (APS) (0.1% m/v; A3678, Millipore Sigma), *N,N,N’,N*’-Tetramethylethylenediamine (TEMED) (0.2% v/v; T9281, Millipore Sigma), and 0.2 µm diameter fluorescent microspheres (0.5% v/v; F8811, ThermoFisher Scientific). This solution was polymerized for 20 min underneath circular glass coverslips (18 mm diameter) to ensure a flat surface. After polymerization, the coverslips were carefully removed, and the gels were rinsed in DI water. Next, sulfo-SANPAH (200 µg/mL; 22589, ThermoFisher Scientific) was conjugated to the gel surface with ultraviolet (UV) light (intensity: 10 mW/cm^2^) for 25 min, followed by DI water rinses. Gels were then incubated in a diluted collagen solution (100 µg/mL; 5226, Advanced Biomatrix) overnight at 4°C. Subsequently, the gels were sterilized under germicidal UV light for 30 min and rinsed with sterile phosphate buffered saline before proceeding with cell culture.

Human synovial fibroblasts (408-05a, Cell Applications) were expanded to ∼85% confluency on tissue culture plastic in synoviocyte growth media (415-500, Millipore Sigma), trypsinized at P5, and labeled with CellTracker fluorescent probe (C34565, ThermoFisher Scientific) for TFM studies. Red fluorescent protein expressing human umbilical vein endothelial cells (RFP-HUVECs) (cAP-0001RFP, Angio-Proteomie) were expanded to 70% confluency on tissue culture plastic in endothelial growth media (EGM-2; CC3162, Lonza) and trypsinized at P5 for TFM studies. Cells were seeded at a low density in each well (1875 cells/cm^2^) to maximize single, non-overlapping cells for analysis. In the co-culture group, synovial fibroblasts and RFP-HUVECs were seeded at a 1:1 ratio, maintaining the same total cell number (half the number of each cell type). Cells attached to the gels within 1 hr (37°C, 5% CO_2_) before administration of IL-1β (100 pg mL^-1^; 20-001B, ThermoFisher Scientific). All groups were cultured in EGM-2 overnight before imaging.

Images were taken with a 20x 0.95 NA water immersion objective at 4x optical zoom (0.2167 µm/px, 1024 x 1024 px) on a Nikon AX Confocal Microscope. All imaging was performed with an environmental chamber (okolab) to control temperature (37°C), humidity (85%), and CO_2_ (5%). The samples were incubated within the microscope chamber for at least 45 min before imaging to reduce z-axis drift during imaging. Within the Nikon software, the ‘perfect focus’ and ‘multi-point imaging’ features were utilized to program the x, y, and z positions for each cell on the gel surface. Images were taken of single, non-overlapping cells before and after lysis. Cells were lysed via administration of 5 drops of 10% m/v sodium dodecyl sulfate (SDS) (436143, Millipore Sigma) into the well.

TFM analysis (stack alignment, particle image velocimetry, and Fourier transform traction cytometry) was performed using a previously developed plugin suite for FIJI (*40*). For FTTC analysis, the Poisson’s ratio of the PA gels was set to 0.45 and a regularization parameter of 1e^-9^ was used, as in (*41*). A custom MATLAB script then inputs the bead displacement and traction stress arrays, and output single cell metrics (i.e., total traction force (nN), max traction stress (Pa)) (Script 2). For coculture gels, the synovial fibroblasts (CellTracker Deep Red) and RFP-HUVECs were separately analyzed.

### Microfluidic device design, assembly, and functionalization

Custom 3D-printed master molds were designed in Autodesk Fusion 360. These 3D-printed molds replaced the previous need for multilayer photolithography to generate masters of the same resolution (*25*, *42*). The new ‘top’ and ‘bottom’ molds were printed using stereolithography (SLA), and Accura 5530 resin (PC-Like Translucent Advanced High Temp resin, Normal Resolution, Natural Finish) with thermal curing (Protolabs). To form the microfluidic devices using these masters, polydimethylsiloxane (PDMS) (2065622, Ellsworth Adhesives) was polymerized in both the top and bottom molds at 60°C overnight. Next, the PDMS layers were demolded, cleaned, aligned, and a PDMS rod (0.337 µm diameter, extracted from 23-gauge needle post-polymerizing) was inserted between the layers. The bi-layered PDMS devices were bonded to glass well-bottom dishes with oxygen plasma (PE25-JW, Plasma Etch).

The microfluidic devices were sterilized under germicidal UV light for 30 min before proceeding with device functionalization. The PDMS was functionalized to promote extracellular matrix attachment. First, a sterile poly-L-lysine (PLL) solution (0.01%; P4707, Millipore Sigma) was injected into the ECM ports, and left for 1 hr at RT. This solution was then removed, and a dilute glutaraldehyde solution (0.4%; G6257, Millipore Sigma) was injected into the ECM ports, and left for 30 min at RT. Subsequently, the devices were thoroughly rinsed three times with DI water and fully dried via aspiration.

### Device seeding and culture

A neutralized collagen solution (3 mg/mL; 5226, Advanced Biomatrix) was prepared on ice ± human synovial fibroblasts (P4-5, 2 million/mL) (408-05a, Cell Applications). In order, reconstitution buffer, 10x DMEM, 1M sodium hydroxide, and DI water were mixed, as described in (*24*). Next, the high concentration collagen solution was added with a wide-bore pipette tip and mixed. Finally, synovial fibroblasts were added to the mixture in a low volume of synoviote growth media. The solution was rapidly transferred into each ECM port of the microfluidic devices. PBS was injected around the perimeter of each dish to prevent evaporation of the collagen solution during polymerization, and the devices were incubated (37°C, 5% CO_2_) for 55 min to polymerize the collagen gels.

Immediately after collagen polymerization, EGM-2 was added to the side ports of each device. The PDMS rods were extracted, allowing perfusion of the media through the hollow lumens. The lumens were flushed twice with media before injecting human umbilical vein endothelial cells (HUVECs) (C2516A, Lonza) into each channel (16k HUVECs/µL). The devices were placed in the incubator and rotated in 90° intervals every 10 min for 80 min to promote even cell attachment. After rotating the devices, the cell solution was removed to eliminate dead, non-adherent cells, and fresh EGM-2 was pipetted into each channel. Lastly, PBS was added around the perimeter of each well to prevent evaporation of the media from the channels.

24 hr after HUVEC seeding, the lumens were washed 5x with EGM-2. Next, EGM-2 ± 100 pg mL^-1^ IL-1β was injected into the channels. Media (EGM-2 ± 100 pg mL^-1^ IL-1β) was changed once daily for up to 8 days.

### Immunofluorescence and imaging

To visualize lumen structure and endothelial inflammation, at the 8-day time point, immunofluorescence staining was performed (VE-cadherin, ICAM-1, DAPI). The lumens were rinsed with PBS, and fixed with 4% v/v paraformaldehyde (15710, Electron Microscopy Sciences) for 15 min at 37°C. The lumens were then rinsed 3x with PBS, and permeabilized with 0.1% v/v Triton-X (A16046.AE, ThermoFisher Scientific) for 15 min at 37°C. Next, the samples were rinsed 3x with PBS and blocked with 5% m/v bovine serum albumin (BSA) (A3311, Millipore Sigma) for 1 hr at RT with agitation. After blocking, the primary antibody solution was added, containing rabbit monoclonal VE-cadherin antibody (1:300 dilution; 2500S, Cell Signaling Technology) and mouse monoclonal ICAM-1/CD54 antibody (1:500 dilution; sc-18853, Santa Cruz Biotechnology) diluted in 5% BSA.

The samples were incubated in this primary solution at 4°C overnight on a shaker. The next morning, the lumens were washed 3x with PBS before adding the secondary antibody solution. The secondary antibody solution contained: donkey anti-rabbit IgG secondary antibody, Alexa Fluor 647 (1:200 dilution; A32795, ThermoFisher Scientific), donkey anti-mouse IgG secondary antibody, Alexa Fluor 488 (1:200 dilution; ab150105, abcam), and DAPI (3:1000 dilution; 62248, ThermoFisher Scientific), diluted in DI water. Samples were incubated in the secondary solution for 2 hr at RT on a shaker and then rinsed 3x with PBS before imaging.

Images were taken with a 20x 0.95 NA water immersion objective with a 1.25x optical zoom (0.69 µm/px, 1024 x 1024 px) on a Nikon AX Confocal Microscope. A z-stack was acquired for each lumen (n = 5 lumens/group), and max projection images were compiled for the 3 channels (VE-cadherin, ICAM-1, DAPI) in FIJI. The max projection images for the ICAM-1 channel were binarized and quantified (%Area along lumen).

### Vascular permeability assay

To determine the influence of the daily IL-1β treatment and synovial fibroblasts on vascular barrier function, permeability assays were performed. Endothelialized lumens (with RFP-HUVECs) ± synovial fibroblasts (from three separate donors, Table S5), and ± daily IL-1β treatment were fabricated and cultured, as described in *device seeding and culture*. Barrier function studies were completed at the 8-day time point using a Nikon AX Confocal Microscope with an environmental chamber (37°C, 5% CO_2_, 85% humidity). Images were taken at the vessel midplane every 3 s for 8 min with a 2x 0.10 NA air objective with a 4x optical zoom (2.16 µm/px, 1024 x 1024 px). After 1-2 images were taken, fluorescent dextran (10 kDa; FD10S, Millipore Sigma) was carefully pipetted into the lumen and began diffusing out of the vessel.

Image series were opened in FIJI and cropped at the center (w: 600 µm, h: 1000 µm) of the vessel, including and above the vessel. Images from the entire time series were exported as .tif files via the image sequence command. Next, a custom MATLAB script, adapted from (*24*), was used to automate the calculation of diffusive permeability from the images.

### Soluble protein release

Soluble cytokines, chemokines, and cell adhesion molecules were analyzed from collected culture media at the 8-day time point. For this analysis, media from 6 individual vessels was pooled per group. Biological triplicates included data from three synovial fibroblast donors (Table S5). A MagPix Luminex Xmap system (ThermoFisher Scientific) was used with a ProcartaPlex Human Inflammation Panel 20Plex (EPX200-12185-901, ThermoFisher Scientific) according to the manufacturer’s protocol. Given the median fluorescent intensity (MFI) readings for each analyte, a custom MATLAB script fit a 4-parameter logistic regression to the standard data. Using this standard curve, the script converted the MFI readings to concentration values (pg mL^-1^). Both the concentration values, and the fold-changes (log2 transformed, normalized to Endo group) of these concentrations were reported.

### Extracellular matrix remodeling and cell localization

Both the endothelial cells and synovial fibroblasts can dynamically remodel their surroundings, contributing to changes in vascular function. To assess the environment around the engineered lumens at the 8-day time point, a small fraction of FITC-conjugated collagen (8% m/v; C4361, Millipore Sigma) was mixed with the unlabeled collagen at the time of gel polymerization (day 0), while maintaining the same overall 3 mg/mL collagen concentration. Six groups were setup for this experiment. The primary experimental groups were Endo (n = 9 lumens), Endo + IL-1β (n = 9 lumens), Endo + Syn (n = 9 lumens), and Endo + Syn + IL-1β (n = 9 lumens). The two control groups were Syn (n = 9 devices), and Collagen alone (acellular) (n = 8 devices).

Images were taken with a 20x 0.95 NA water immersion objective with a 1.25x optical zoom (0.69 µm/px, 1024 x 1024 px) on a Nikon AX Confocal Microscope. A z-stack (250 µm depth, 5 µm slice thickness) was acquired for each sample (n = 9 lumens/group) starting at the vessel midplane and going down (past the bottom of the lumen). Images were resliced in FIJI to create orthogonal slices, along the length of the lumen. Each vessel was uniformly cropped in the radial direction. Images were converted to an 8-bit format, and uniformly thresholded and de-noised. Lastly, the 3D Objects Counter command (FIJI) was implemented to compute the volume of the collagen layer surrounding each lumen.

Using the same devices, synovial fibroblast localization was assessed. The synovial fibroblasts were stained with cell tracker deep red at the time of cell seeding and RFP-HUVECs were used to endothelialize the lumens, enabling live imaging. The Syn + Endo and Syn + Endo + IL-1β groups were compared to determine whether IL-1β treatment influenced synovial fibroblast localization along the lumens. For image analysis, max projection images of the synovial fibroblast channel were compiled in FIJI. Images were cropped to encompass only the vessel area, thresholded, and binarized. Lastly, the %Area covered by synovial fibroblasts was measured.

### Statistical analyses

To assess grader-to-grader variability for each histological scoring metric, the intraclass correlation (ICC) was calculated using as custom MATLAB script. An ICC score ≥ 0.6 signified sufficient grader-to-grader agreement for that metric. All other statistics were computed using GraphPad Prism. For data involving the comparison of only two groups, an unpaired t-test was performed. For parametric data with > two groups, but only one categorical variable, a one-way ANOVA was run, followed by Tukey’s post-hoc comparisons. For parametric data with β two groups and two categorical variables (i.e., ± synovial fibroblasts, and ± Il-1β treatment), a two-way ANOVA was run followed by multiple comparisons with Fishers LSD test. For all analyses, \**P*< 0.05, \*\**P*< 0.01, \*\*\**P*<0.001, \*\*\*\**P*<0.0001.

## Acknowledgments

The authors would like to thank Dr. Joseph Dragavon and the BioFrontiers Advanced Light Microscopy Core (RRID: SCR_018302) for advice and guidance pertaining to image acquisition. The authors are also grateful to Sung Yeon Kim and Dr. Edward Bonnevie for openly sharing documentation related to the fabrication of polyacrylamide gels for TFM. The authors lastly thank Arkodip Mandal for providing feedback on this work that improved the overall manuscript.

## Author contributions

Conceptualization: HMZ, CRS, LEH, JAB

Methodology: HMZ, DNG, CJC, APD, MDD, ASP, HKW, MW

Investigation: HMZ, DNG, CJC, APD, MDD, ASP, HKW, MW

Visualization: HMZ, DNG Supervision: CRS, LEH, JAB

Writing—original draft: HMZ

Writing—review & editing: HMZ, DNG, CJC, APD, MDD, ASP, HKW, MW, CRS, LEH, JAB

## Data and materials availability

All data are available in the main text or the supplementary materials.

