## Supplement for "Synovial fibroblasts support vascular function in an acute injury-on-a-chip model"

Hannah M. Zlotnick *et al.*

*Jason A. Burdick.

**This PDF file includes:**

Figs. S1 to S6
Tables S1 to S5
Legend for Data S1

**Other Supplementary Materials for this manuscript include the following:**

Data S1

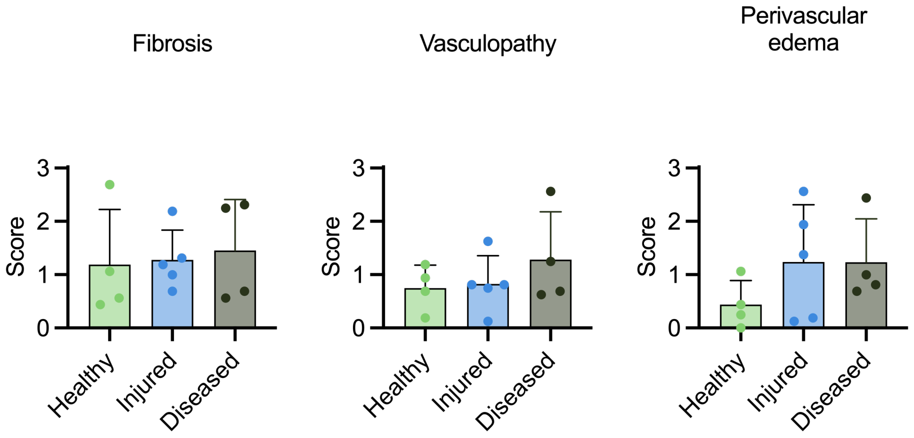


**Figure S1. Semi-quantitative histological scoring of fibrosis, vasculopathy, and perivascular edema in human synovia.** Histological scoring of human knee synovia (Healthy: n=4; Injured: n=5; Diseased: n=4) stained with hematoxylin and eosin. For each scoring criterion, scores ranged from 0 (physiologic) to 3 (pathophysiologic). Injured: meniscus injury. Diseased: total knee replacement. n = 4 trained, blinded scorers. Data plotted as the average score for each tissue section by 4 scorers and bars represent mean + std dev. ANOVA with Tukey post-hoc comparisons, no significance between groups for fibrosis, vasculopathy, or perivascular edema.

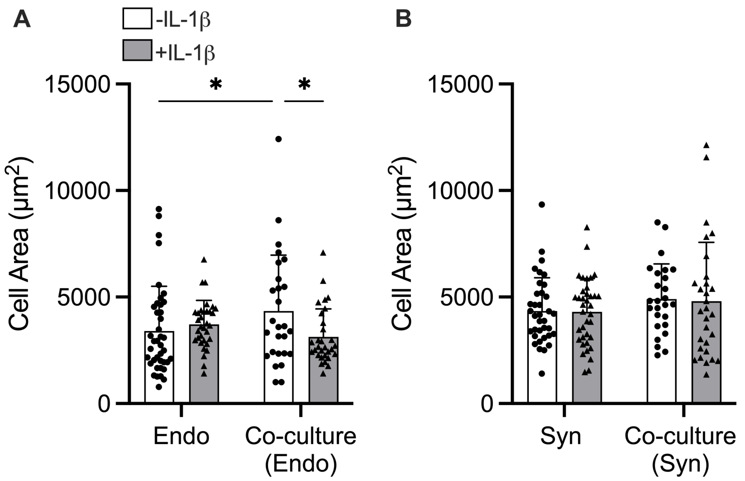


**Figure S2. Area of endothelial cells and synovial fibroblasts cultured alone or in co-culture on polyacrylamide (PA) gels.** (A) Endothelial cell area when cultured in isolation and co-culture (with synovial fibroblasts). White bars = cultured in EGM-2, grey bars = cultured in EGM-2 + 100 pg/mL IL-1β. n = 27-40 cells/group. 2-way ANOVA with Fisher's LSD test. **P* <0.05. (B) Synovial fibroblast cell area when cultured in isolation and co-culture (with endothelial cells). White bars = cultured in EGM-2, grey bars = cultured in EGM-2 + 100 pg/mL IL-1β. n = 26-37 cells/group. 2-way ANOVA with Fisher's LSD test. No significance between groups. Bars represent = mean + std dev.

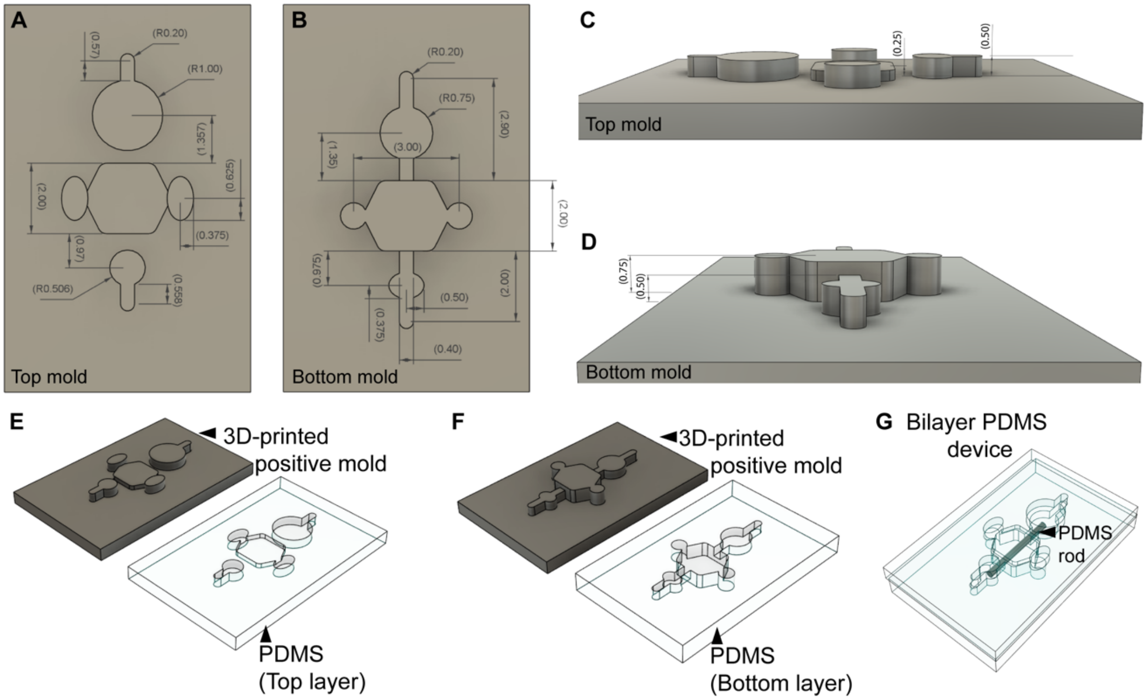


**Figure S3. 3D printed mold designs for use in microfluidic devices and PDMS casting.** (A, B) Aerial view of top and bottom layer molds with dimensions (in mm). (C, D) Side view of top and bottom layer molds with dimensions (in mm). (E, F) 3D printed positive molds (a single unit), and the PDMS negative used for devices. (G) Bi-layered PDMS device with inner PDMS rod used during collagen polymerization (rod removal creates open channel within the collagen gel).

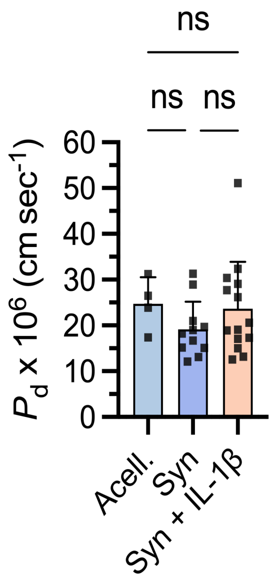


**Figure S4. Diffusive permeability ($P_d$) of FITC-Dextran (10 kDa) in control groups without endothelialized lumens.** Acell: 3 mg/mL collagen, cultured in EGM-2, no cells (only collagen). Syn: Synovial fibroblasts (2 million/mL) in collagen gel, no endothelialized lumen, cultured in EGM-2. Syn + IL-1β: Synovial fibroblasts (2 million/mL) in collagen gel, no endothelialized lumen, cultured in EGM-2 + 100 pg/mL IL-1β. n = 4 devices for Acell, n = 11-14 devices for Syn, and Syn + IL-1β. No significance between groups (ns), 1-way ANOVA with Tukey post-hoc comparisons. Bars represent mean + std dev.

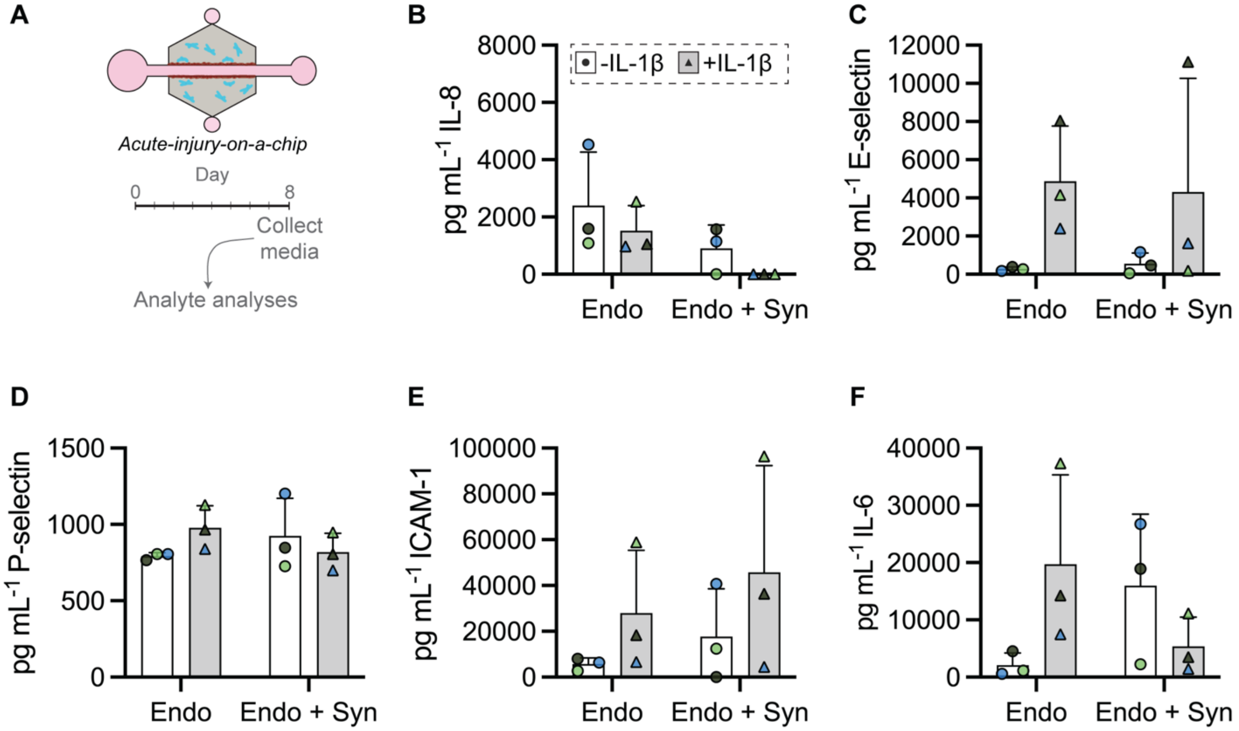


**Figure S5. Soluble protein release from acute injury-on-a-chip model system.** Endo: endothelialized lumen cultured in EGM-2. Endo + IL-1β: endothelialized lumen cultured in EGM-2 + IL-1β. Endo + Syn: endothelialized lumen surrounded by human synovial fibroblasts (in the collagen gel) cultured in EGM-2. Endo + Syn + IL-1β: endothelialized lumen surrounded by human synovial fibroblasts (in the collagen gel) cultured in EGM-2 + IL-1β. (A) Overview of media collection. (B-F) Concentrations (pg/mL) of secreted proteins from multiplex inflammation panel without statistical significance (assessed via 2-way ANOVA with Fisher's LSD test). Colored replicates (n = 6 lumens pooled/group/donor). Bars represent overall mean + std dev.

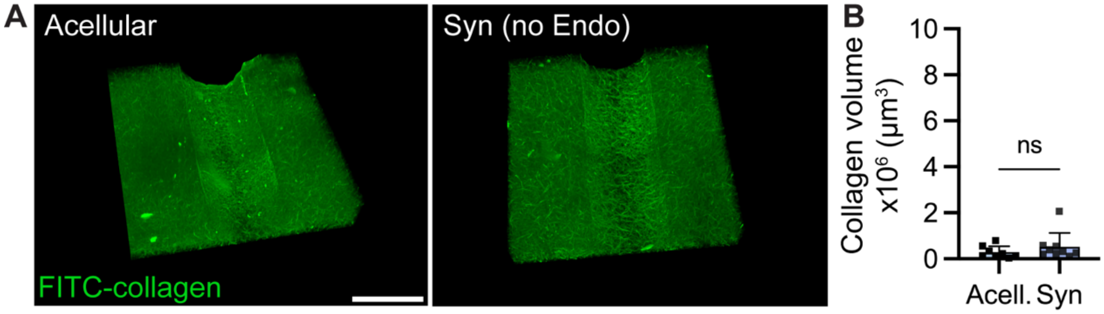


**Figure S6. Extracellular matrix (collagen) within microfluidic devices is uniform in the absence of endothelial cells.** (A) Acellular and synovial fibroblast (Syn) controls, both without endothelial cells. Scale bar = 400 µm. (B) Volumetric analysis of FITC-collagen along the length of the lumen. There is minimal, if any, remodeling / artifact observed. n = 8-9 lumens/group. Unpaired t-test. ns: not significant, defined by $P<0.05$. Bars represent mean + std dev.

**Table S1.**
Demographics for organ donor patients. Collins Grade 0: Normal; Grade 1: Limited disruption of the articular surface; Grade 2: Fibrillation of cartilage with fissures, ± small osteophytes; Grade 3: Extensive fibrillation and fissuring; 4: >30% of the cartilage eroded down to the subchondral bone, gross geometric changes + osteophytes. M: male. F: female.

| Organ donor | Age (years) | Sex | Collins Grade |
|---|---|---|---|
| 1 | 24 | M | 0 |
| 2 | 73 | M | 1 |
| 3 | 38 | F | 1 |
| 4 | 66 | M | 3 |
| 5 | 62 | F | 3 |

**Table S2.**
Demographics for meniscal arthroscopy patients. KL: Kellen and Lawrence classification system for radiographic scoring. 0: no OA, 4: severe OA. M: male. F: female. NR: not reported, score < 3.

| Meniscal arthroscopy (Meniscus injury) | Age (years) | Sex | KL-classification |
|---|---|---|---|
| 1 | 56 | F | NR |
| 2 | 79 | F | 2 |
| 3 | 77 | F | 2 |
| 4 | 36 | M | 1 |
| 5 | 72 | F | 2 |

**Table S3.**
Demographics for total knee arthroplasty (TKA) patients. KL: Kellen and Lawrence classification system for radiographic scoring. 0: no OA, 4: severe OA. F: female. NR: not reported.

| Total knee arthroplasty (TKA) | Age | Sex | KL-classification |
|---|---|---|---|
| 1 | 69 | F | 4 |
| 2 | NR | NR | 3/4 |
| 3 | NR | NR | 3/4 |
| 4 | NR | NR | 3/4 |

**Table S4.**
Intraclass correlation (ICC) values (two-way mixed-effects model for consistency, single measurement) for histological scoring parameters.

| Parameter | Intraclass correlation (ICC) value |
|---|---|
| Infiltrates | 0.61 |
| Vascularity | 0.74 |
| Lining Hyperplasia | 0.68 |
| Fibrosis | 0.65 |
| Vasculopathy | 0.58 |
| Perivascular Edema | 0.73 |

**Table S5.**
Donor information for human cells. M: Male. F: Female. NB: Newborn. B: Black. W: White. A: Asian. NR: Not reported.

| Human Primary Cells | Vendor | Cat # | Lot/Batch # | Age | Sex | Race |
|---|---|---|---|---|---|---|
| Synovial Fibroblasts | Cell Applications | 408-05a | 3606 | 67 | M | B |
| Synovial Fibroblasts | Cell Applications | 408-05a | 3617 | 58 | M | W |
| Synovial Fibroblasts | Cell Applications | 408-05a | 3823 | 65 | M | W |
| Human umbilical vein endothelial cells (Pooled) | Lonza | C2519A | 23TL086130 | NB | M/F mixed | A/W |
| RFP-Human umbilical vein endothelial cells | Angio-Proteomie | cAP-0001RFP | 2022050402 | NB | Pooled, NR | Pooled, NR |

**Data S1. (separate file)**
Raw data and statistical analysis for all main text and supplementary figures.
